# Adhesion of *Crithidia fasciculata* promotes a rapid change in developmental fate driven by cAMP signaling

**DOI:** 10.1101/2022.10.06.511084

**Authors:** Shane Denecke, Madeline F. Malfara, Kelly R. Hodges, Nikki A. Holmes, Andre R. Williams, Julia H. Gallagher-Teske, Julia M. Pascarella, Abigail M. Daniels, Geert Jan Sterk, Rob Leurs, Gordon Ruthel, Rachel Hoang, Megan L. Povelones, Michael Povelones

## Abstract

Kinetoplastids are single-celled parasites responsible for human and animal disease. For the vast majority of kinetoplastids, colonization of an insect host is required for transmission. Stable attachment to insect tissues via the single flagellum coincides with differentiation and morphological changes. Although this process is essential for the generation of infectious forms, the molecular mechanism driving differentiation following adherence is not well understood. To study this process, we elaborate upon an in vitro model in which flagellated swimming cells of the kinetoplastid *Crithidia fasciculata* rapidly differentiate following adhesion to artificial substrates. Manipulation of culture parameters revealed that growth phase and time had a strong influence on the proportion of adherent cells. Live imaging of cells transitioning from swimming to an attached cell fate show parasites undergoing a defined sequence of events including an initial adhesion near the base of the flagellum, immediately followed by flagellar shortening, cell rounding, and the formation of a hemidesmosome-like structure between the tip of the shortened flagellum and the substrate. We have also assayed the role of the cyclic AMP (cAMP) signaling pathway in differentiation of *C. fasciculata*. Pharmacological inhibition of cAMP phosphodiesterases eliminated the ability of swimming cells to attach without affecting their growth rate. Further, treatment with inhibitor did not affect the growth rate of established attached cells, indicating its effect is limited to a critical window of time during the early stages of adhesion. Finally, in swimming parasites we have shown that a receptor adenylate cyclase localizes to the distal portion of the flagellum. In attached cells it is absent from the shortened flagellum and instead localizes to the cell body. Similarly, a putative phosphodiesterase, found along the length of the flagellum, also relocalizes to the cell body in attached parasites. These data suggest that in *C. fasciculata* cAMP signaling is required for adherence, that cAMP flux in the flagellum of swimming cells is spatially restricted, and that signaling domains may be reorganized during differentiation and attachment. These studies contribute to our understanding of the flagellum as a multi-functional organelle integrating processes related to motility, signaling, attachment, and differentiation and further develop *C. fasciculata* as a model kinetoplastid.

**Author Summary:** Parasites from the order Kinetoplastida are transmitted by insects and cause diseases such as Leishmaniasis, Chagas disease, and Human African Trypanosomiasis. A key aspect of the life cycle of these parasites is their ability to adhere to surfaces within the insect and subsequently differentiate into forms that are transmissable to their next host. Here, we explore the molecular mechanisms underpinning both adherence and differentiation using the mosquito parasite *Crithidia fasciculata*. This organism is closely related to pathogenic species but grows to high densities in culture and adheres robustly in vitro. We identified factors affecting the rate of adherence and defined morphological stages of this process including rapid shortening of the cell’s single flagellum. Using electron microscopy, we confirmed that adhesion to artificial substrates results in the formation of a filament-rich adhesive plaque at the tip of the flagellum that resembles the attachment to insect tissue. We also demonstrate the key role played by the cyclic AMP signaling pathway during adherence and differentiation. Specifically, pharmacological inhibition of enzymes that degrade cyclic AMP completely blocks adherence, and tagging of several proteins in this pathway show that their localization along the flagellum changes following differentiation to the attached form. These data provide insights into processes critical for all kinetoplastid life cycles.

## Introduction

Organisms from the protozoan order Kinetoplastida impose a significant disease burden especially in low-income countries [1]. *Trypanosoma brucei, Trypanosoma cruzi*, and *Leishmania* spp. are the etiological agents for African trypanosomiasis, Chagas disease, and Leishmaniasis, respectively. These parasites also cause important veterinary diseases, while still others impact insects critical for pollination. A deeper understanding of the biology of these organisms can provide the basis for rational interventions to limit disease burden. Additionally, kinetoplastid parasites are a compelling model for basic research, representing early-branching eukaryotes that are divergent from more well-studied organisms such as insects, worms, and vertebrates.

Kinetoplastids primarily parasitize insects with only a handful of species using insects as a vector to infect other organisms. In insects, kinetoplastids spend part of their life cycle adhered to the apical surface of an epithelial lining. Adherence, also referred to as adhesion or attachment in the literature, often precedes a developmental transition with distinct morphological and molecular changes. For example, in the kissing bug *Rhodnius prolixus, T. cruzi* parasites bind to the perimicrovillar membrane in the posterior midgut, followed by the cuticular lining of the hindgut, before being excreted by the insect as infectious metacyclics [2, 3]. Adherence is necessary not only to maintain the parasites in the gut of their host, but to trigger differentiation to the infectious form. Similarly, *T. brucei* adheres to salivary gland epithelium and *Leishmania* spp. adhere to the midgut and the stomodeal valve. In each case, adherence precedes or is concurrent with differentiation to the final developmental form in the insect. Despite the importance of adherence in the kinetoplastid insect life cycle, the relative inaccessibility of these parasite stages in the insect, and the lack of a robust in vitro model to generate them, means that little is known about adherence mechanisms and the signal transduction pathways that may connect adherence to developmental progression.

*Crithidia fasciculata* is a monoxenous parasite of mosquitoes that developmentally transitions from a swimming (nectomonad) cell with a long flagellum to an attached (haptomonad) cell with a dramatically shortened flagellum. The tip of this flagellum is attached to the chitinous lining of the hindgut epithelium via a hemidesmosome-like adhesive plaque. A morphologically similar plaque is found in the adhered forms of other kinetoplastid species [4]. Remarkably, for *C. fasciculata* this developmental switch from a swimming to an attached cell is readily produced in vitro by adhesion to a variety of substrates, including Millipore filters [5], culture debris [6], and plastic dishes [7, 8]. Taking advantage of the ability to produce large numbers of swimming and attached cells in culture, our recent transcriptomic analysis identified genes encoding putative cyclic AMP (cAMP) signaling components that were differentially regulated in the different forms of *C. fasciculata*. This led to the hypothesis that cAMP signaling may be involved in cell fate transitions in these organisms [7].

Despite its importance, much remains unknown about cAMP signaling in kinetoplastids [9]. The cAMP second messenger is created from ATP by adenylate cyclases (ACs), many of which are transmembrane receptor adenylate cyclases (RACs) that probably act as receptors for unidentified ligands. The RAC family of ACs is dramatically expanded in *T. brucei*, in which they localize to the flagellum, either along the length or at the distal tip [10]. In many systems protein kinase A (PKA) acts downstream of cAMP, but in kinetoplastids, [11, 12], PKA is independent cAMP [13]. Instead, several cAMP response proteins (CARPs) are thought to mediate downstream events. Phosphodiesterases (PDEs) terminate the signal by degrading cAMP to 5’-AMP. In *T. brucei*, some of these PDEs are localized to the flagellum, while others are found in the cell body [14]. It has been proposed that PDEs act to limit the diffusion of cAMP and create spatially restricted signaling microdomains [15]. This paradigm has also been demonstrated in other eukaryotic systems [16].

Using a robust and quantitative adherence assay and imaging of live and fixed cells, we have defined the process of adherence and the conditions that promote it in vitro. In addition, we have used pharmacological inhibitors to probe the role of cAMP signaling during the transition to an attached state. Finally, we show the localization of putative components of the *C. fasciculata* cAMP signaling pathway to the flagellum in swimming cells and show striking changes in the distribution of these proteins upon flagellar shortening. These findings support the use of *C. fasciculata* as a model to understand the spatial regulation of signal transduction in the context of adherence and differentiation of kinetoplastid parasites.

## Results

### Culture conditions impact in vitro adherence of *C. fasciculata*

To probe the mechanism of *C. fasciculata* adherence, we used an in vitro assay to determine which conditions impact the adherence rate. Swimming cells, grown on a shaker, were transferred to fresh culture plates and incubated without shaking to allow cells to adhere. Plates were washed to remove non-adherent cells and then the number of single adhered cells in representative fields were quantitated (Fig 1A). Over the next 24 hours, these single cells divide to form attached rosettes. Cells in a rosette can undergo division to produce either two attached cells or two swimming cells that can leave the rosette. Some proportion of these newly-produced swimming cells can also encounter the dish and become attached. In our standard assay, swimming cells were again washed away at 24 hours and rosettes with four or more cells in representative fields were quantitated.

**Figure 1.**
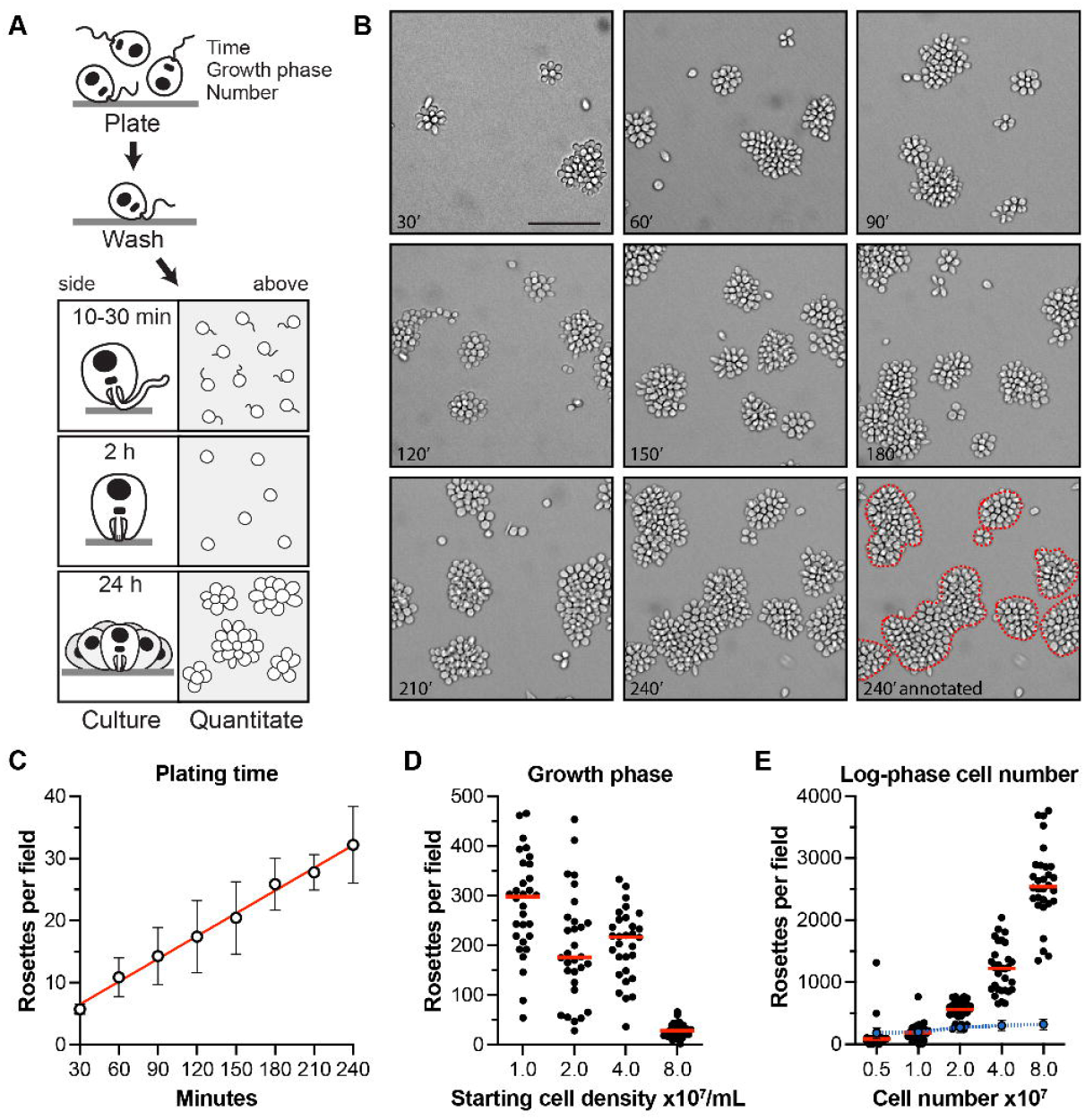
An adherence assay reveals features of *C. fasciculata* attachment. **A)** Schematic showing steps of the assay, including the variables tested. **B)** Representative images showing attached rosettes produced by varying the initial plating time. The dashed red lines were drawn to indicate what was counted manually as individual rosettes, clusters containing 4 or more cells. **C)** Quantitation of the experiment shown in B. This experiment was performed in triplicate. Error bars show standard error. **D)** The effect of initial cell concentration on adherence. 10^7^ cells/mL from cultures in different phase of growth, were adhered for each sample. Quantitation of a representative experiment of three replicates is shown with median values indicated by red lines. **E)** Rosettes resulting from adherence of different numbers of cells, all derived from a mid-log phase culture (10^7^ cells/mL). A representative experiment of three replicates is shown. Red lines indicate median values, and the blue dotted line represents the proportion of plated cells that were able to adhere.

By varying the length of plating time prior to washing, we found that the length of this incubation step was directly proportional to the number of attached parasites, with the longest time we tested, 240 minutes, producing the greatest number of rosettes (Fig 1 B,C). Consistent with a previous report [17], we also found that the growth phase of the cultured cells had a strong effect on the rate of adherence. Cells from an early log phase culture (10^7^ cells/mL) adhered and differentiated at a higher frequency compared with the same number of cells taken from a stationary phase culture (10^8^ cells/mL; Fig 1D). In addition to having a reduced rate of cell division, stationary phase cells are morphologically distinct from cells in exponentially growing cultures, with the latter being rounder and shorter (Figure S1). Increasing the total number of cells used in the adherence assay, provided those cells were taken from a culture in log-phase, proportionally increased the total number of attached cells but had relatively little impact on the percentage of cells that adhered during the 2 hour incubation (Fig 1E). These data suggest that in vitro adherence of *C. fasciculata* happens at a constant rate and is inversely correlated with the culture density before the assay begins but not while adherence occurs.

### Mechanism of attachment of *C. fasciculata* in vitro

To examine the stages of adherence and differentiation of *C. fasciculata* more precisely, we used time-lapse microscopy to follow parasites in environmentally controlled conditions. Using flagellar shortening and cell rounding as morphological markers for differentiation, we observed that these events were always preceded by an initial adhesion event (Videos S1,S2). During this event, cells first adhere to the plate via their anterior end, near the flagellar pocket, and then begin to retract their flagellum. As this happens, the cells become rounder as they undergo their first cellular division as adherent cells. In certain cases, cell division occurs prior to the full retraction of the flagellum, but this was rare. Parasites sometimes adhered in pairs, a phenomenon that has been reported previously [18]. In this case, one cell of the pair will typically detach and swim away (Video S2). From these experiments, it was also possible to directly measure the rate of flagellar shortening, revealing that the flagellum retracts at a relatively constant rate until it is no longer visible at around 2 hours (Fig 2).

**Figure 2.**
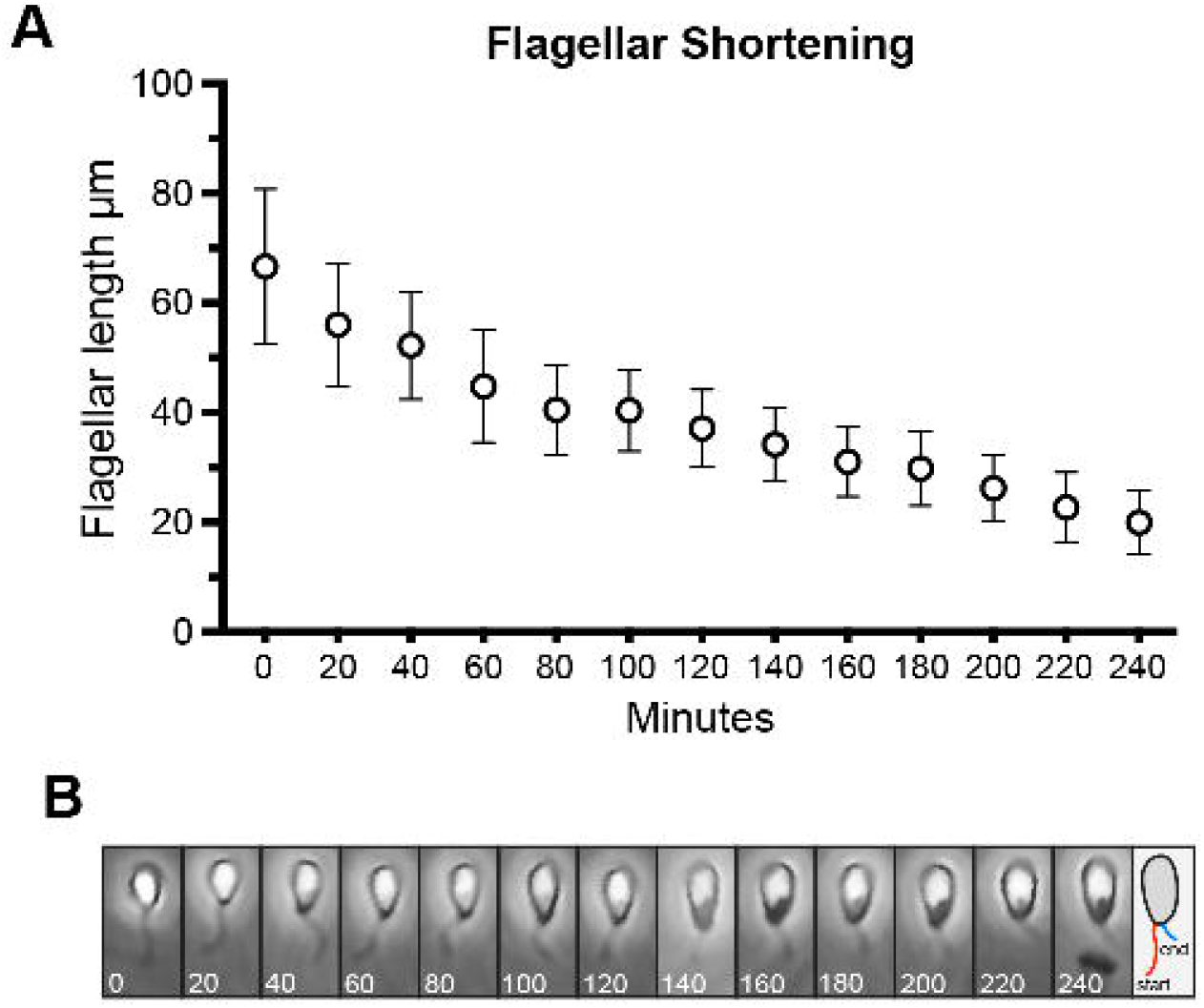
Flagellar shortening occurs at a relatively constant rate following an initial adhesion event. **A)** Following 5 minutes of plating and subsequent washing, cells were observed at the indicated times and the length of the flagellum was measured in 20 representative cells (different cells were measured at each time point). **B)** An example of a single cell undergoing flagellar shortening. The numbers indicate time in minutes. The schematic on the right shows a representation of the length of the flagellum at the beginning (red) and end (blue) of the time course.

While parasites initially adhere near the base of their flagellum, stably attached cells are connected to the substrate via the distal tip of the shortened flagellum. Previous reports on *C. fasciculata* and other kinetoplastids have described plaques at the site of attachment that ultrastructurally resemble hemidesmosomes [6]. Since most imaging of the adhesive structure was performed on parasites adhered to the insect, we used transmission electron microscopy (TEM) to determine if hemidesmosomes were present in attached cells where they contact the culture plate. Consistent with prior findings [8], a cross-section of rosettes showed an osmophilic, filamentous plaque at the distal tip of the shortened flagellum adjacent to the substrate (Fig 3 A-B). When we detached adherent cells from culture plates by scraping prior to fixation, the hemidesmosome appeared intact, indicating that attachment mediated by this structure can be physically disrupted (Fig 3C). Fibrous, desmosome-like structures appear to connect the proximal part of the flagellum to the cell body at the flagellar pocket, features which have also been noted in prior studies and which are probably components of the flagellar attachment zone (FAZ) (Fig 3A-C) [6, 19, 20]. When cells were sectioned *en face* near the attachment site with the culture plate, we observed that the distal portion of the flagellum containing the hemidesmosome was somewhat enlarged and extended beyond the termination point of flagellar axoneme (Fig 3D). In these sections, hemidesmosomes are circular and the degree to which they are stained by uranyl acetate is likely a function of how close to the culture plate they were sectioned (Fig 3D,E). By contrast, in swimming cells, we only found FAZ-like structures connecting the flagellum to the pocket but did not observe hemidesmosomes at the distal end of the flagellum (Fig 3F). A notable feature of both attached cells and scraped attached cells was an extracellular network of filamentous material attached to the plasma membrane of the cell body (Fig 3G and 3A-D). In swimming cells, similar material was sometimes observed in the flagellar pocket itself, but not on the cell surface (Fig 3F).

**Figure 3.**
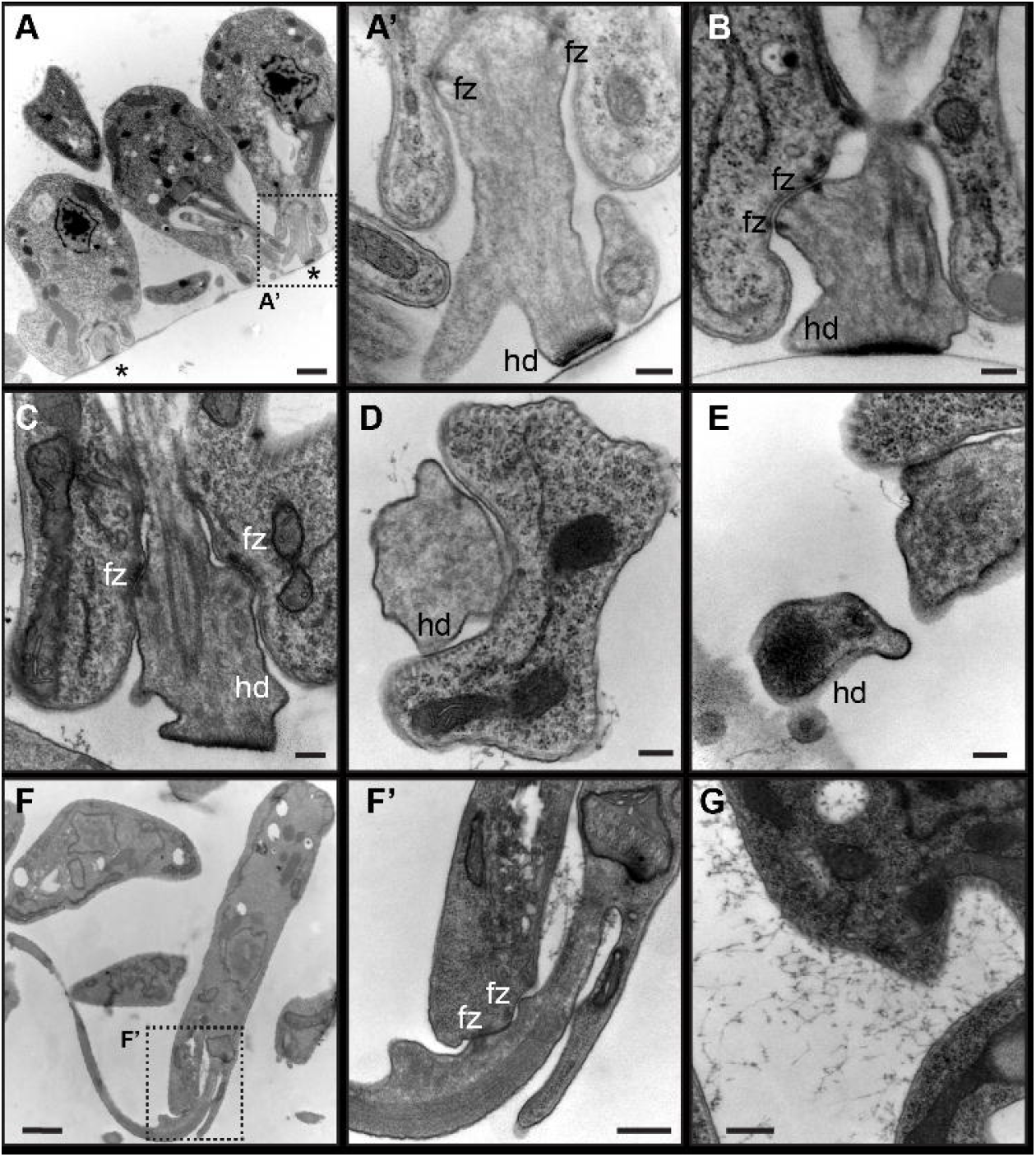
Transmission electron microscopy reveals structural differences between attached and swimming cells. **A)** Cross section through *C. fasciculata* parasites adhered to tissue culture plastic (Fig S2, yellow plane). Asterisks indicate hemidesmosomes at the distal tip of the shortened flagella. Scale bar is 1 μm. The dotted box indicates the region enlarged in panel A’. **A’)** Close up of a hemidesmosome (hd) from one of the cells in panel A. Desmosome-like FAZ structures, connecting the flagellum to the cell body, are also observed (fz). Scale bar is 200 nm. **B)** and **C)** Additional examples of hemidesmosomes (hd) and desmosome-like FAZ (fz). The cell in C) was scraped from the dish prior to fixation. Scale bar is 200 nm. **D)** Sectioning *en face* shows a region near the end of the flagellum in an adherent cell (Fig S2, blue planes). The flagellar membrane is enlarged and no flagellar axoneme can be observed. Instead, diffuse filaments probably comprising the hemidesmosome are seen. Scale bar is 200 nm. **E)** Another example of a hemidesmosome sectioned *en face*. Scale bar is 200 nm. **F)** Section of a swimming cell shows the flagellum extending from the flagellar pocket. Dotted box indicates the region enlarged in F’. Scale bar is 1 μm. **F’)** Higher magnification view of the flagellar pocket of a swimming cell. FAZ structures link the flagellar membrane to the flagellar pocket membrane, but no hemidesmosome is seen. The slight bulge in the flagellum where it exits the pocket may be a physical correlate to the site of initial adhesion. Filamentous material can be observed in the flagellar pocket of the swimming cell but was never observed on the surface. In contrast, the surface of attached cells, **G**) was covered with this material. Some of the same material is also visible in panels A, B, C, D, and E. Scale bar F’ and G is 400 nm.

### Pharmacological Inhibition of PDEs blocks adherence without affecting growth

Our prior work showed that cAMP signaling components are upregulated in swimming *C. fasciculata* relative to attached parasites [7]. To test whether manipulating the cAMP pathway could influence the ability of *C. fasciculata* to adhere to plastic in vitro, we used previously validated compounds that are known to inhibit cAMP PDEs in the related parasite, *T. brucei* [21]. As PDEs terminate the signal by degrading the cAMP second messenger, inhibition of these enzymes has been shown to increase the amount of cAMP in the parasite [22]. The first compound we tested is known as NPD-001 or Compound A (CpdA) (hereafter NPD-001). The addition of 10 μM of NPD-001 to the adherence assay completely blocked parasite attachment to the plate, resulting in no single adhered cells after 2 hours of incubation and no visible rosettes after 24 hours of growth (Fig 4A). Interestingly, a related compound, NPD-008, had only a minor effect on adhesion, slightly reducing the number of single adhered cells after 2 hours of incubation (Fig 4A). To determine if the effect of NPD-001 on attachment was dose-dependent, we performed an adherence assay with both 1 μM and 0.1 μM of the compound. Although the effect of 1 μM was comparable to that of 10 μM, 0.1 μM had a moderate effect on adhesion, suggesting a dose response. The same concentrations of a related compound, NPD-226, also significantly reduced the number of adhered cells both at 2 hours and at 24 hours at all three concentrations compared to vehicle-treated controls (Fig 4B). A fourth compound, NPD-055, had a minor effect on adherence at the highest concentration (10 μM, Fig S3).

**Figure 4.**
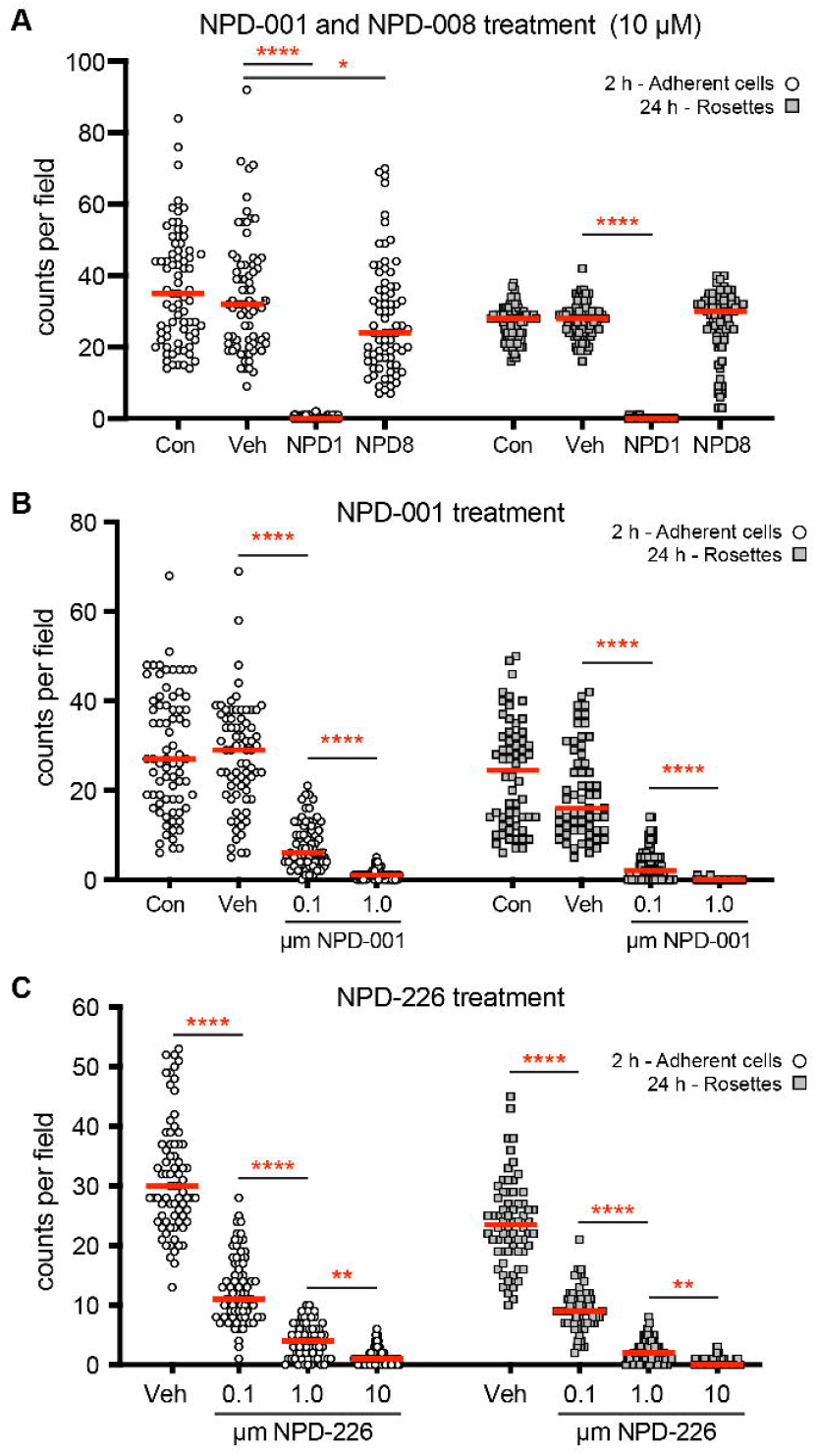
Phosphodiesterase inhibitors block *C. fasciculata* adherence. **A**) Adherence assays performed in the presence of 10 μM of either NPD-001 (NPD1) or NPD-008 (NPD8). The number of single adhered cells at 2 hours (open circles) and the number of rosettes (≥ 4 cells, gray squares) at 24 hours were counted and compared to those from a vehicle-treated control sample (Veh) and an untreated sample (Con). Treatments were present in the media both during the 2-hour plating time and the subsequent 24 hours of culture. All comparisons were made. Asterisks indicate Kruskal-Wallis test with Dunn’s correction P<0.0001 (****) and P<0.05 (*) **B**) Adhesion assays comparing vehicle and untreated controls to assays performed in the presence of different concentrations of NPD-001 or **C**) NPD-226. The number of single adhered cells at 2 hours (open circles) and the number of rosettes (≥ 4 cells, gray squares) at 24 hours were counted. All comparisons were made, but only the groups that are significantly different from the adjacent treatment are shown. Asterisks indicate Kruskal-Wallis test with Dunn’s correction P<0.0001 (****), and P<0.01 (**).

To confirm that the adherence-blocking activity of the PDE inhibitors was not due to cell toxicity, we measured growth rates of swimming cultures of *C. fasciculata* (maintained on a shaker) in the presence of NPD-001. Addition of this compound to the media did not affect the growth rate of swimming cells compared to vehicle- or untreated control cells (Fig 5A). Furthermore, addition of NPD-001 to cells that were already differentiated to the attached state did not affect their attachment or their doubling time over the course of 16 hours (Fig 5B). In this experiment, we did observe new adherence of swimming cells produced from the rosettes in the vehicle-treated control. Consistent with our previous finding (Fig 4A), although new swimming cells were produced from the rosettes, in NPD-001 treated cultures, they failed to adhere.

**Figure 5.**
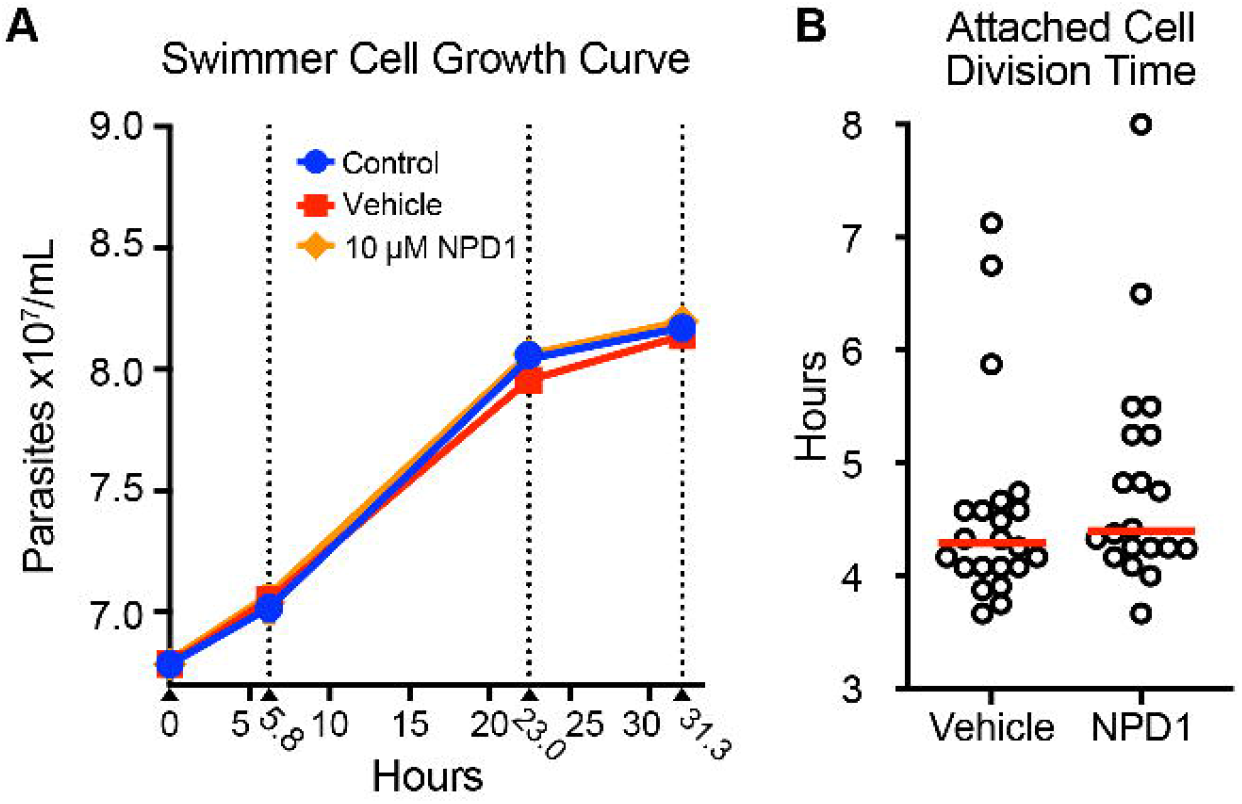
NPD-001 does not affect the growth rate of *C. fasciculata*. **A)** Graph of *C. fasciculata* growth in shaking cultures assayed over approximately 31 h in standard medium (Control; blue circles), medium supplemented containing 10 μM NPD-001 in vehicle (NPD1; orange diamonds), or medium containing vehicle only (Vehicle, red squares). **B**) Time encoded movies were used to measure the doubling time of established attached cultures after the addition of NPD-001 in vehicle, or in vehicle alone. The red line indicates the mean. No significant difference was noted between the doubling times.

### cAMP signaling proteins are localized to the flagellum

In other kinetoplastids, components of cAMP signaling pathways are localized to flagellar subdomains, indicating that this organelle may be involved in spatially restricting the distribution of this second messenger [10, 14]. To see if this feature is also present in *C. fasciculata*, we localized a putative receptor adenylate cyclase (*Cf*RAC1) and a putative cAMP phosphodiesterase (*Cf*PDEA) by fusing a GFP protein to the C-terminus of each open reading frame and introducing these constructs into parasites as episomes (Fig S4). In swimming *C. fasciculata*, *Cf*PDEA::GFP was found along the length of the flagellum, often appearing as two parallel lines of puncta (Fig 6). The extent of *Cf*PDEA::GFP is somewhat diminished along the portion of the flagellum in the flagellar pocket. *Cf*RAC1::GFP was found in similar puncta, but was restricted to the distal third of the flagellum in swimming parasites (Fig 6, Fig S5). We compared both of these localization patterns to another cell line in which a YFP sequence was fused to one of the endogenous copies of the *Cf*PF16 gene (Fig S4), a component of the central pair of the flagellar axoneme [23]. *Cf*PF16::YFP was uniformly distributed along the length of the flagellum in a single continuous line without obvious puncta. No fluorescent signal was detected in parental CfC1 cells (Fig S6).

**Figure 6.**
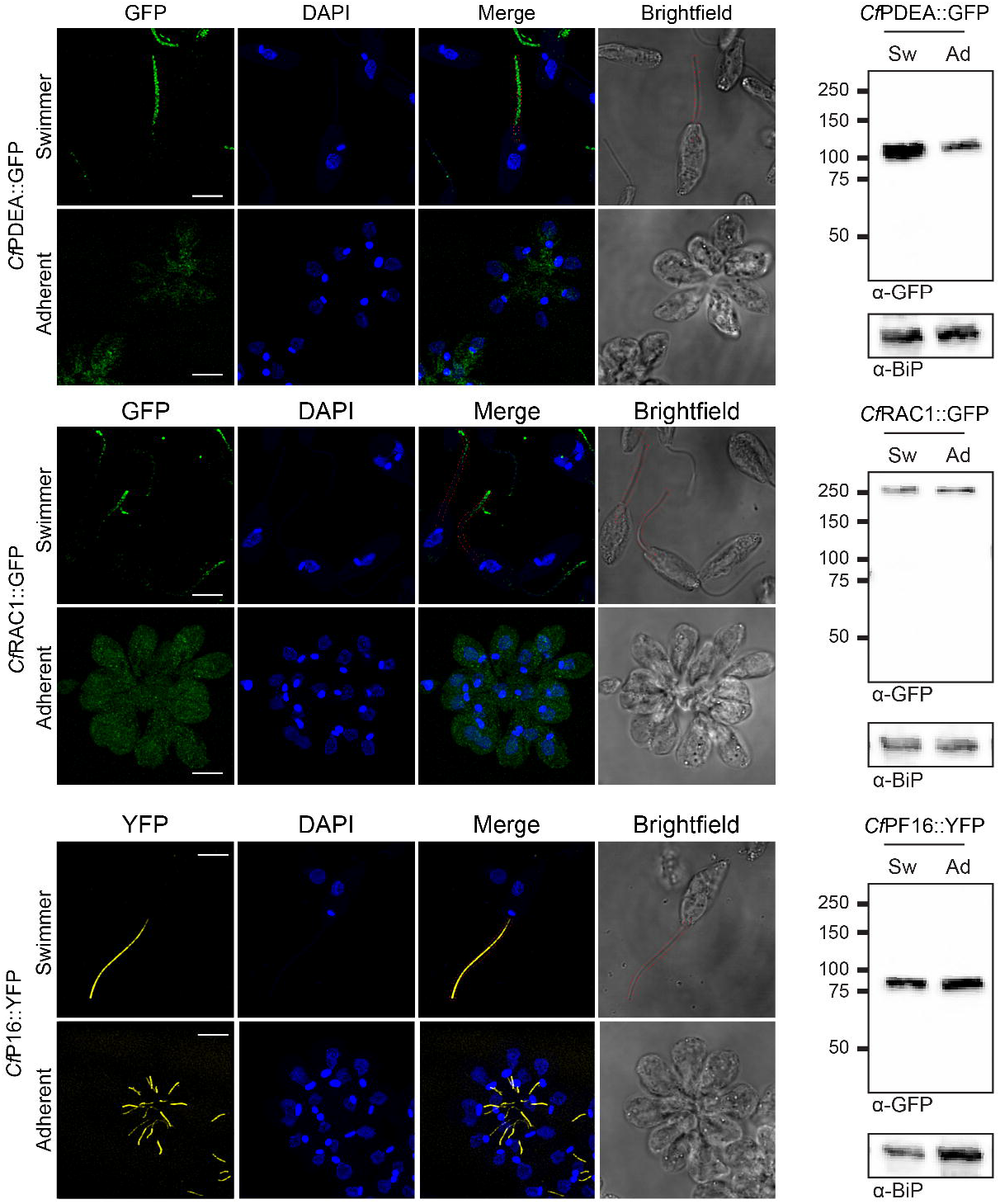
Components of cAMP signaling localize to different domains of the flagellum in *C. fasciculata* swimmers. Deconvolved confocal images of GFP and YFP fusion proteins and DAPI staining of nuclear and kDNA of fixed swimmer and attached cell cultures. A Z-stack of the entire cells and rosettes were captured. Fluorescent images are shown individually and merged. A brightfield image for each is included for reference and was used to generate the red dashed outline of the flagellum in the merged and brightfield images. An overlay of fluorescent channels on the corresponding brightfield image also shows the extent of GFP or YFP signal with respect to the length of the flagellum (Fig S4). The scale bars are 5 μm. Western blots of extracts of swimmers (Sw) and attached (Ad) cells of each line are included supporting the presence of the correct size GFP- or YFP-fusion protein (Fig S3) in both swimmer and attached cells regardless of their localization patterns. Equivalent loading was confirmed by re-probing blots with an anti-BiP antibody. Parental Cf-C1 cells lack GFP signal in swimmers and adherent cells as well as by western analysis (Fig S5).

We then followed these localization patterns as cells transitioned from the swimming to the attached form. As expected, *Cf*PF16::YFP could still be detected in the shortened flagellum which, as supported by our EM studies, retains the central pair. However, *Cf*RAC1::GFP was no longer localized to the flagellum and was instead distributed across the entire cell body. A similar redistribution was also observed in attached cells expressing *Cf*PDEA::GFP (Fig 6). To compare expression levels of these proteins in attached and swimming *C. fasciculata*, we performed western blotting using an anti-GFP antibody. Consistent with our microscopy results, *Cf*PDEA::GFP and *Cf*RAC1::GFP are readily detected in both swimming and attached parasites despite their different subcellular localizations (Fig 6). No fluorescent or anti-GFP western signal was observed in the parental cells (Fig S6).

## Discussion

Adherence and differentiation are both critical for the successful completion of kinetoplastid life cycles. While signal transduction pathways driving differentiation have been described in multiple species, virtually nothing is known about mechanisms of attachment and how this process is related to critical developmental events in the insect vector. Numerous studies have found important roles for cAMP-based signaling in differentiation and response to host environments, yet many questions remain unanswered in this area as well, including why kinetoplastids have expanded their repertoire of RACs, what signals are sensed, and how downstream signals are transduced. In addition, the significance of signaling subdomains in the flagellum is not understood. *C. fasciculata* provides an opportunity to dissect the relationship between flagellar signaling and adherence in the context of differentiation. We expect that, with some variations, this connection exists in all kinetoplastids. Our ability to follow adherence and differentiation in live cells, coupled with a quantitative assay for adherence and expanding tools for molecular genetics, opens a wide range of avenues for future exploration.

### The mechanism of *C. fasciculata* attachment in vitro

The adherence of *C. fasciculata* and other kinetoplastids to substrates in vitro has been noted previously [6, 8, 17, 24–27]. Having developed a reproducible, quantitative assay for this phenomenon, we decided to explore factors that would increase or decrease adherence. Consistent with previous work, we found that adherence of *C. fasciculata* to artificial surfaces results in the formation of a hemidesmosome-like structure at the point of attachment at the tip of the shortened flagellum. Due to the similarity of this attachment structure to that formed in vivo [28, 29], we can use the in vitro system to determine the conditions required for its formation. As was reported previously [17], we have also found that cultures in stationary phase are less likely to adhere compared to log phase cultures, with a large drop off in adherence for parasites growing between 4 to 8×10^7^ cells/mL. *C. fasciculata* parasites in stationary phase were shown to have a more elongated cell morphology compared to log phase cells, and previous studies have demonstrated differential protein abundance between these cell states possibly linked to their nutritional status [30]. Such nutrient deprivation has been shown to trigger adherence of *T. cruzi* epimastigotes after which the parasites differentiate to the infectious metacyclic form [31]. In our assays, this effect seems to result from an inherent difference between parasites in log-versus stationary-phase, as plates seeded with large numbers of log phase cells can form nearly confluent layers, suggesting that there is no upper limit to density in this assay. As adherence is linearly related to the number of plated log-phase cells, we also conclude that it is not cooperative, and that each parasite adheres independently to the substrate. This is supported by our TEM, live-cell, and fluorescence imaging studies which confirm that rosettes are formed by cell division of individually adhered cells, which are each attached to the surface through their flagellum. In this case, the number of attached parasites should be limited by substrate surface area. Consistently, previous studies on *Leishmania* spp. have noted that scratching the surface, thereby increasing the available surface area, increases parasite adherence [27], although this did not appear to trigger differentiation. It should be noted that the term “rosette” has also been used to describe an observed tendency of some *Leishmania* spp. parasites to adhere to each other via their flagella [32–35]. These rosettes are not adhered and are not formed through cell division, but instead result from clustering of parasites. Thus far we have not seen this phenomenon in our *C. fasciculata* cultures.

We have also used this assay to probe the mechanism of adherence. *C. fasciculata* parasites initially contact the dish via the anterior end of the cell body. We call this initial event “adhesion” which is rapidly followed by flagellar shortening, cell rounding, and formation of the hemidesmosome. We call this fully differentiated state “attached” reflecting the stable attachment between the tip of the flagellum and the substrate presumably facilitated by the hemidesmosome-like structure and accompanied by changes in gene expression. In this study, we have found that even at log phase the percentage of *C. fasciculata* parasites that stably attach to the dish is low and increases linearly with time. Furthermore, our live-cell imaging studies show that not every cell that encounters the plate undergoes differentiation to the attached form. These aborted adhesion events suggest that only a subset of cells can make the transition at any one time, which may reflect stochastic variation, timing within the cell cycle, or other factors. For example, it was shown previously that a subset of stationary-phase *C. fasciculata* fail to agglutinate after treatment with peanut lectin, indicating possible variations in cell surface molecules [30]. These glycoproteins and glycoconjugates may connect to the membrane by glycosylphosphatidylinositol (GPI) anchors, which are found commonly among kinetoplastids. For example, it has been suggested that differences in cell surface lipophosphoglycan (LPG) between promastigotes and metacyclics in *L. major* may underpin the ability of the former to adhere to the midgut of their sandfly vector [36, 37]. These findings are particularly interesting in the context of the TEM data reported here, which show a lattice-like filamentous molecule at the cell surface only in adherent cells. Similar material was detected in the flagellar pocket of swimming cells, suggesting that it may be synthesized and secreted by this form, but not retained on the surface. Determining the identity of this molecule will allow for exploration of possible roles in adhesion or protection in the insect gut, as well as comparisons with other kinetoplastid species.

The predictable regularity of *C. fasciculata* attachment suggests a developmental program resulting in a distinct cell state. Flagellar shortening occurs at a constant rate and seems triggered by the initial adhesion event. Our TEM images show that the shortened flagellum of attached *C. fasciculata* are distinct from that of *Leishmania* amastigotes in that the *C. fasciculata* axoneme retains the central pair of microtubules even at the tip. In addition to the formation of hemidesmosome filaments, the *C. fasciculata* flagellar membrane at the point of attachment is also expanded, as has been noted for haptomonads of other species [24, 26, 38]. From these data and our experiments with NPD-001/CpdA, it appears that the final attached state, once established, is relatively stable. As in other studies, we have found that the hemidesmosome is extremely resistant to detergents and enzymatic digestion [39] but can be physically disrupted by scraping [40].

### The role of cyclic AMP in *C. fasciculata* adhesion and differentiation

Strikingly, we found that inhibition of PDEs with NPD-001 completely blocked attachment. As treatment of stably attached *C. fasciculata* parasites with NPD-001 did not cause the cells to detach nor did it prevent division of attached cells, we hypothesize that elevated cAMP resulting from PDE inhibition disrupts the dynamic transition from initial adhesion to stably attached. An alternative interpretation is that elevated cAMP impacts the swimming cell population by dramatically reducing the subset of cells that are competent to adhere, meaning that these cells would not even form initial adhesions. Our findings contribute to a large body of work showing that signaling via the cAMP pathway plays a critical role in developmental transitions in kinetoplastid parasites [9, 41], including the epimastigote to metacyclic transition in *T. cruzi* [42] and the transitions to stumpy [42] and procyclic form *T. brucei* [43]. However, in *T. brucei* there is evidence that differentiation may be triggered by degradation products of cAMP, rather than cAMP itself [44]. In addition to differentiation, *T. brucei* requires cAMP signaling for social motility, a process by which parasites generate and sense pH gradients and migrate collectively through the tissues of their insect host, eventually attaching to the salivary gland epithelium and differentiating to infectious metacyclics [12, 45, 46]. Treatment with CpdA/NPD-001 blocks social motility in vitro [45], and genetic knockout of the *T. brucei* PDEB1 gene or other signaling components impairs or blocks developmental progression through the fly [12, 46]. Due to the impact on social motility, the effect of these manipulations on *T. brucei* attachment cannot be readily assessed.

Variations in flagellar length are a common feature of kinetoplastid life cycles [47], which may impact the distribution of flagellar cAMP signaling proteins. We have found that GFP fusions of a putative RAC and PDE (*Cf*RAC1::GFP and *Cf*PDEA::GFP) both localize to the flagellum in swimming *C. fasciculata*, with *Cf*RAC1::GFP being enriched in the distal tip. Both *Cf*PDEA::GFP and *Cf*RAC1::GFP display punctate localization patterns compared to *Cf*PF16::YFP. While *Cf*RAC1 has a predicted transmembrane domain, *Cf*PDEA does not. However, in *T. brucei*, the phosphodiesterase TbPDEB 1 is an integral component of the flagellum that is resistant to detergent extraction [15]. Lipid analysis of the flagellar membrane has suggested lipid-raft like membrane heterogeneity, which could also lead to uneven localization of our tagged proteins [48]. Therefore, anchoring of *Cf*RAC1 and *Cf*PDEA to the flagellar membrane or structures within the flagellum could explain the puncta that we observe. In addition, the parallel lines of signal found in both tagged cell lines is reminiscent of intraflagellar transport trains, which associate with two sets of microtubule doublets on either side of the axoneme [49]. The different distributions of these flagellar proteins, *Cf*RAC1 to the tip and *Cf*PDEA to the entire length, is also consistent with observations in other organisms, in which specific amino acid sequences are required for correct localization to flagellar subdomains [10]. However, how these domains are established and the mechanisms for protein targeting are not yet clear.

Both tagged proteins are no longer enriched in the shortened flagellum of attached parasites, indicating differential localization in the two developmental forms of *C. fasciculata*. As expected, *Cf*PF16::YFP was retained in the shortened flagellum, as this structure still has its central pair of microtubules. It is unclear from these studies if resident signaling proteins are relocalized from the flagellum to the cell body during transition from swimming to attached, or whether newly synthesized signaling proteins in attached cells fail to enter the flagellum, perhaps due to alterations in the machinery for flagellar protein sorting. Fluorescence time-course imaging of differentiating cells should help reveal the dynamic localization patterns of cAMP signaling components, which would convey critical information about this process.

Our data suggest that cAMP signaling acts during a relatively narrow window to facilitate the transition between initial adhesion and stable attachment. As illustrated in Figure 7, our model is that the physical contact with the surface during adhesion initiates flagellar shortening leading to local alterations in cAMP flux which then trigger a developmental program including changes in transcript abundance and formation of the attachment plaque. This would mean that the spatial restriction of RACs, PDEs, and other signaling proteins conveys important statiotemporal information that is vital for successful developmental transitions. *C. fasciculata* is an attractive system in which to tease out these processes, and future studies will explore the connection between flagellar dynamics, signaling, adherence, and differentiation in this important group of parasites.

**Figure 7.**
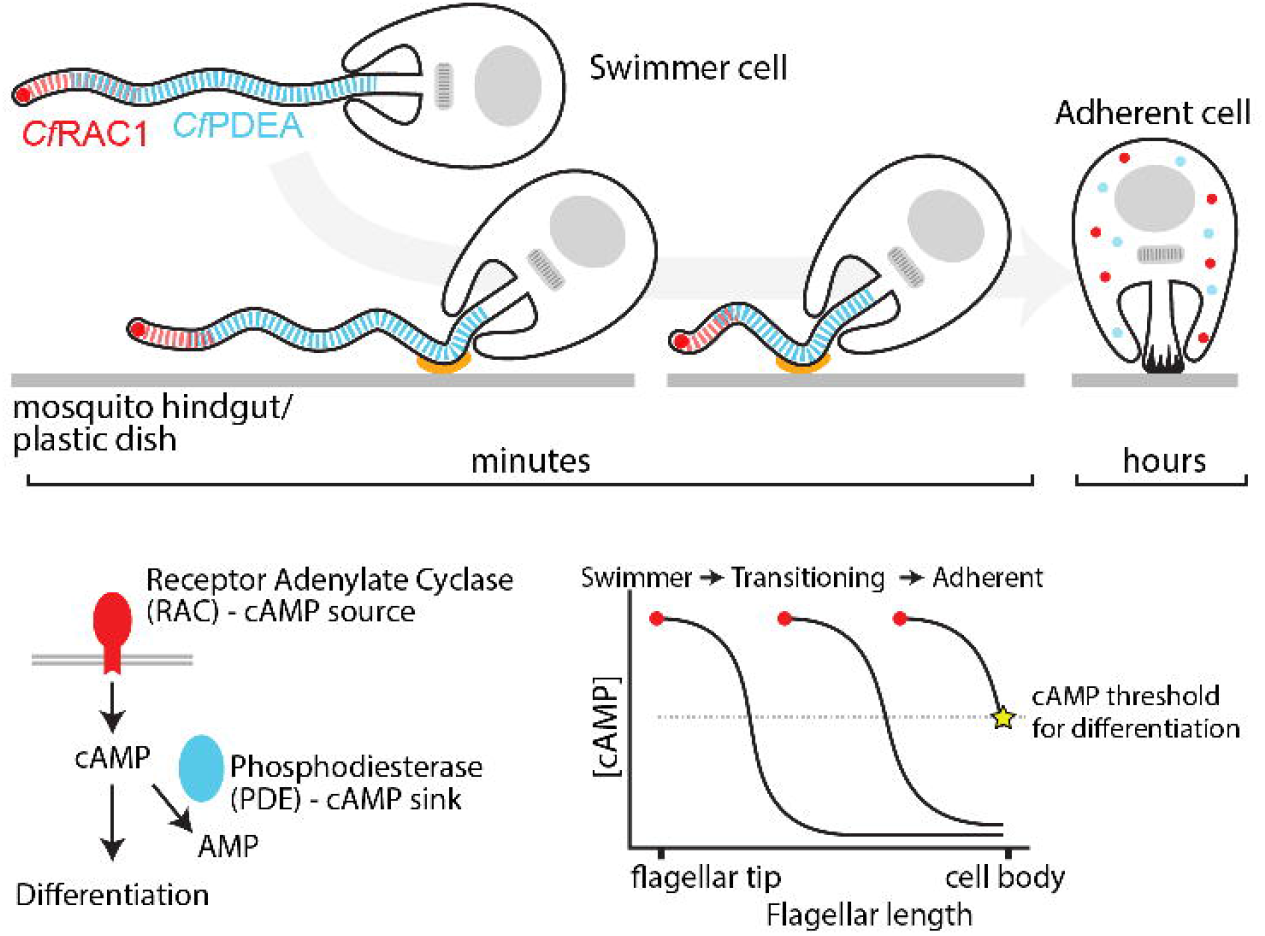
Model of the developmental transition from swimmer to adherent cell. The localizations of *Cf*RAC1 (red), and *Cf*PDEA (blue) are indicated. Minutes following adhesion via the region of the flagellum near the cell body (orange), the process of flagellar shortening begins. Flagellar shortening brings *Cf*RAC1 closer to the cell body. Eventually the source of cAMP is close enough to the cell and the sink for cAMP (*Cf*PDEA) is reduced such that cAMP levels rise above a critical threshold in the cell body (shown on the graph by a star). This promotes differentiation to the adherent cell fate characterized cytoplasmic localization of *Cf*RAC1 and *Cf*PDEA by robust changes in gene regulation and formation of a hemidesmosome.

## Materials and Methods

### *Crithidia* maintenance

The Cf-C1 strain of *C. fasciculata* was used for all experiments in this study. This strain was originally obtained from Dr. Stephen Beverley and was grown in complete medium (BHI) consisting of Brain Heart Infusion medium (Sigma) supplemented with 20 μg/mL hemin (Sigma). Cells were maintained at 28 °C in non-vented culture flasks on a rocker and were maintained at a density between 10^5^ and 10^8^ cells/mL. Swimming cells were counted on a hemocytometer following fixation in 0.3% formalin.

### Plasmids and cell lines

The construct for C-terminal YFP tagging of the *C. fasciculata* putative axoneme central apparatus protein CFAC1_170028000, named *Cf*PF16 for its shared homology to *L. mexicana* PF16 [23], was created using a fusion PCR approach described previously [50]. The YFP and neomycin resistance genes were amplified from pLENTv2eYFPNeo by PCR with forward primer pLENTeYFP_f and reverse primer pLENTNeo_r (Table 1). 500 bp of homology to the C-terminus of *Cf*PF16 was amplified from *C. fasciculata* genomic DNA using forward primer PF16cterm500_f and reverse primer PF16cterm_r. 500 bp of homology to the 3’ UTR of *Cf*PF16 was amplified using PF16utr_f and PF16utr500_r. All three fragments were purified by PCR clean-up or gel extraction and 20 ng of each fragment was combined in a fusion PCR reaction using PF16_500bpnest_f and PF16_500bpnest_r. The resulting construct, PF16eYFPNeo, was gel extracted and ethanol precipitated. To create a *C. fasciculata* CfPF16::YFP cell line, transfection was performed with 3.5 μg of PF16eYFPNeo introduced into 10^8^ cells using a Human T-cell kit and IIb Nucleofector set to program X-001 (Lonza). Other transfections described in this work used the same conditions except with *Tb*-BSF buffer (90 mM sodium phosphate, 5 mM potassium chloride, 0.15 mM calcium chloride, 50 mM HEPES, pH 7.3) [51]. Immediately following transfection, cells were transferred to 30 mL of BHI without selecting drug and diluted 10-fold. For each dilution, 1 mL was plated into wells of a 24-well plate and cultured overnight without shaking. After approximately 18 hours of recovery, 1 mL of BHI containing neomycin (G418) was added to each well, for a final selecting drug concentration of 25 μg/mL. Resistant clones came up in 7-14 days and were fixed and screened for eYFP fluorescence using wide-field microscopy. The resulting clonal cell line CfPF16YFP_A6 was used for these studies and was validated by linking PCR across the gene with primer PF16ORF_f binding inside the gene and reverse primer PF16UTR_r binding in the 3’UTR (outside of the area used for integration). To create the *C. fasciculata Cf*RAC1::GFP and CfPDEA::GFP cell lines, the entire open reading frame of *Cf*RAC1 (CFAC1_090006500) or CfPDEA (CFAC1_140020300) were amplified from *C. fasciculata* genomic DNA and cloned into the pNUSGFPcH vector [52] using NdeI and KpnI or NdeI and BglII restriction enzyme sites, respectively. Following confirmation by sequencing (Eurofins Genomics), 10 μg of the final plasmid was introduced into *C. fasciculata* as described above. Transfected cells were selected using BHI containing 80 μg/mL hygromycin. After 7 days, selecting drug concentration was increased to 200 μg/mL and cells were screened by fluorescence microscopy.

### Adherence assay

Establishment of cultures of swimming and adherent *C. fasciculata* in vitro were adapted from our previous study [7]. Briefly, cells from a rocking culture of swimmers were added to 60 mm non-treated tissue culture plates (VWR) in 2 mL of BHI. For some experiments, cells were spun down in tubes at 800 rcf for 5 minutes and resuspended in BHI. The cell suspension was allowed to sit in the 60 mm plates undisturbed at 28 °C for the indicated amounts of time during which adherence takes place (2 hours for the standard assay). Following this incubation, each plate was washed 5 times with PBS before addition of fresh BHI media. When assays were performed in 24-well plates, the same protocol was followed but the wash volume was 2 mL. Other assay parameters were tested systematically and described in the results.

Quantitation of adherence was performed through imaging on a Zeiss Primovert Digital Microscope. Following wash steps, plates were imaged by taking either 25 frames at 40x magnification or 10-25 frames at 20x magnification. Freshly adhered cells (singlets) were counted either manually or, when large numbers of adhered cells were present, by Fiji script [23]. Rosettes were counted manually as clusters of 4 or more cells at 24 hours.

### Transmission electron microscopy and Giemsa staining

Swimming and attached *C. fasciculata* cells were processed and imaged using transmission electron microscopy (TEM) performed in the Electron Microscopy Resource Lab (Perelman School of Medicine, University of Pennsylvania). For swimming cells, 10^8^ mid-log-phase cells were concentrated by centrifugation and resuspended in TEM fix (0.1 M sodium cacodylate buffer, pH 7.4, containing 2.5% glutaraldehyde, and 2.0% paraformaldehyde). For attached cells, rosette cultures in 60 mm dishes were established for 24 hours as described above. 5 mL of TEM fix was added to the plate at room temperature. A second 24-hour attached sample was generated by scraping cells off the plate with a disposable plastic scraper, concentrating by centrifugation, and placing in TEM fix. All samples were fixed initially at room temperature for 2 hours and then placed at 4 °C overnight. After subsequent buffer washes, samples were post-fixed in 2.0% osmium tetroxide with 1.5% K3Fe(CN)6 for 1 hour at room temperature and rinsed in water prior to *en bloc* staining with 2% uranyl acetate. After dehydration through a graded ethanol series, the tissue was infiltrated and embedded in EMbed-812 (Electron Microscopy Sciences). Thin sections were stained with uranyl acetate and SATO lead and examined with a JEOL 1010 electron microscope fitted with a Hamamatsu digital camera and AMT Advantage NanoSprint500 software. Images of cells cut in cross section and *en face* were captured (Fig S2). Morphological changes between mid-log phase and stationary phase cells were obtained by performing a cytologic cytocentrifuge preparation and a standard Wright-Giemsa stain and imaging at 100x.

### High content imaging

High content imaging was performed using an ImageXpress Micro 4-High Content imaging device (Molecular Devices) at 40x and 60x using standard 24 (CellTreat) and black welled (Vision 4tituder) 24 well plates respectively. To measure rosette growth, cells were allowed to adhere to standard plates for 2 hours followed by 5 washes with 2 mL of PBS and resuspension in fresh BHI media. Adhesion videos were obtained using black welled plates by adding 4 x 10^5^ cells to 500 μL of BHI media and allowing adhesion for 2 hours prior to imaging at 1.25 min intervals.

### Confocal microscopy and image processing

For fluorescence and confocal microscopy, 5 x 10^6^ swimming cells from mid-log cultures were concentrated by centrifugation at 800 rcf for 5 minutes, washed once in PBS, then resuspended in PBS at a final concentration of 10^7^ cells/mL. 500 μL of this suspension was then pipetted onto coverslips pretreated with 0.01% poly-L-lysine. After allowing cells to attach for 20 minutes at room temperature, coverslips were washed twice with PBS, followed by fixation in 4% paraformaldehyde for 15 minutes. Coverslips were then washed again followed by permeabilization with 0.1% Triton X-100 in PBS for 5 minutes. Coverslips were washed in PBS prior to staining with 0.2 μg/mL DAPI in PBS for 5 minutes. After a final PBS wash, coverslips were mounted on a glass slide using Vectashield mounting medium (Vector Laboratories). For fluorescence microscopy of attached cells, a standard adherence assay was conducted, as described above, on 1.5 thickness glass-bottom dishes pre-coated with poly-L-lysine (MatTek). After 24 hours of adherent growth, the final BHI wash contained 16% formaldehyde added to a final concentration of 4%. The plate was processed from this point as described above for the swimmer cell slides. The slides and plates were analyzed by confocal microscopy using a Leica SP5-II with a 100x objective. A Z-series was acquired for GFP/YFP, DAPI and brightfield. Following image acquisition, deconvolution analysis was performed on the fluorescent images using Huygens Essential deconvolution software (version 17.04.1p2 64b; SVI). Brightfield images are single confocal sections with a flatfield correction applied. All images of parental, *Cf*PDEA::GFP, and *Cf*RAC1::GFP were acquired with the same microscope settings. Equivalent linear adjustments were made to the brightness of each image using Photoshop Version 23.5.1 (Adobe).

### Western blot analysis

For swimming cultures, samples were generated from 2×10^6^ log-phase cells. For adhered cultures, an equivalent number of cells were grown on 60 mm dishes in stationary culture for 24 hours and then dislodged using a cell scraper and counted by hemacytometer. Both swimming and dislodged adherent cells were transferred to microcentrifuge tubes and collected by centrifugation, washed once with PBS, then resuspended in Laemmli SDS-PAGE sample buffer at a concentration of 1×10^5^ cell equivalents/μl. Samples boiled for 5 min at 95 °C and centrifuged briefly. Cell equivalents of ~1.5×10^6^ cells per lane were separated by 10% SDS-PAGE along with the Precision Plus Protein Kaleidoscope standard (BioRad). The fractionated proteins were transferred to a PVDF membrane. The membrane was then incubated in blocking buffer (PBS containing 5% milk and 0.2% Tween-20) for 1 hour at room temperature. A mouse anti-GFP antibody (Roche) was used as the primary antibody at a 1:500 dilution in blocking buffer overnight at 4 °C. Horseradish peroxidase-conjugated goat anti-mouse IgG in blocking buffer was used as the secondary antibody (1:5000). After washing, Enhanced Chemiluminescent Substrate (BioRad) was used to develop the blots. For the loading control, the blot was reprobed with a rabbit anti-*Tb*BiP antibody (a gift of Jay Bangs) at a 1:5000 dilution in blocking buffer, followed by washing and probing with an anti-rabbit-HRP secondary antibody (1:5000). Images were taken by an Azure biosystems c600 using chemiluminescence settings.

### Pharmacological inhibition

Pharmacological inhibitors were dissolved in DMSO. For vehicle controls, an amount of DMSO equal to that of the most concentrated sample was added. The compounds were described previously: NPD-001 [22], NPD-008 [53], NPD-055 [54], and NPD-226 [55].

### Statistical analyses and data visualization

Prism Version 9.4.1 (458) (GraphPad Software) was used to generate all plots and for all statistical tests. Plots were assembled into figures using Illustrator Version 26.5 (Adobe).

## Supporting information

Striking Image

## Acknowledgments

We thank Biao Zuo for preparing TEM samples and assistance with image acquisition and Koranda Walsh and the PennVet Clinical Pathology lab for processing the Giemsa staining samples. We thank Emmanuel Tetaud, Sam Dean, Eva Gluenz, and Tom Beneke for plasmids and advice. Thanks to Stephen Beverley for providing the Cf-C1 wild-type *C. fasciculata* cell line and who, along with Wesley Warren, Chad Tomlinson, and Peter Myler, made the unpublished genome sequence available on TriTrypDB. We thank Jay Bangs for providing the anti-BiP antibody and Lindsay Bair, Laura Anastor-Walters, and Nancy Peltier for their technical assistance. We thank Jesse Quatse for assistance with Photoshop. Finally, we acknowledge the PennVet Imaging Core for providing access to instrumentation and analysis software used in live-cell and confocal imaging and image processing.

---

**Figure S1.** Density dependent changes to *C. fasciculata* morphology. **A**) Giemsa-stained samples of cells at log-phase (10^7^ cells/mL) or past log-phase growth (10^8^ cells/mL). Scale bar is 10 μm. **B**) Plot of individual measured cell lengths in micrometers. Cells were measured along the anterior-posterior axis of the cell body excluding the flagellum. The data are normally distributed (Shapiro-Wilk test). The red line indicates the mean and standard deviation. An unpaired t test indicates the cell length distributions are significantly different with a P value of 0.005.

**Figure S2.** Illustration of sectioning planes used for transmission electron microscopy. An adherent cell shown in cross section (yellow plane; sections in Fig 3A-C, and F-G). Sections cut *en face* at different distances from the culture plate (blue planes). Depending on the position of the *en face* section relative to the culture plate, the osmiophilic hemidesmosome-like structure will appear darker (Fig 3E) or lighter (Fig 3D) and other cell features, such as the cell cytoplasm and mitochondrion may be visible (Fig 3D).

**Figure S3.** Phosphodiesterase inhibitor NPD-055 partially reduces *C. fasciculata* adherence. Adherence assays performed in the presence in the presence of different amounts of NPD-055 compared to vehicle-only treated controls. The number of single adhered cells at 2 hours (open circles) and the number of rosettes (≥ 4 cells, gray squares) at 24 hours were counted and compared to those from a vehicle-treated control sample (Veh). Treatments were present in the media both during the 2-hour plating time and the subsequent 24 hours of culture. Data are not normally distributed. Red lines indicate the median. All comparisons were made, but only the groups that are significantly different from the adjacent treatment are shown. Asterisks indicate Kruskal-Wallis test with Dunn’s correction P<0.0001 (****), P<0.001 (***), and P<0.05 (*).

**Figure S4.** Schematic diagram of tagged proteins used for localization studies. Open reading frames encoding *Cf*PDEA (CFAC1_140020300) and *Cf*RAC1 (CFAC1_090006500) were cloned without their C-terminal stop codon into a vector resulting in an in-frame fusion with the coding sequence of GFP (green rectangle). The resulting constructs were transfected into *C. fasciculata* and lines episomally expressing the tagged proteins were selected using hygromycin. A single allele of *Cf*PF16 (CFAC1_170028000) was fused in-frame at its C-terminus with YFP (yellow rectangle) by homologous recombination followed by neomycin selection. The number of amino acids (aa) for each fusion protein is provided. Domains and their relative positions indicated in the different constructs are 3’5’-cyclic nucleotide phosphodiesterase, catalytic domain (PDEase; IPR002073), Periplasmic binding protein-like I (PBPL; IPR028082), Nucleotide cyclase (AC; IPR029787), Armadillo-type fold (Armadillo-fold; IPR016024), signal peptide for secretion (SP), and transmembrane domain (TM). Further details are provided in the Materials and Methods.

**Figure S5.** Overlay images show the extent of fluorescent signal along the swimming cell flagellum. Overlay of the brightfield and fluorescent images of parental Cf-C1, *Cf*PDEA::GFP, *Cf*RAC1::GFP, and *Cf*PF16::YFP from Figure 6. The fluorescence of *Cf*PDEA::GFP reaches the cell body and weakly inside, similar, but less uniform as *Cf*PF16::YFP. In contrast, *Cf*RAC1::GFP is more restricted to the distal part of the flagellum. Parental cells lack GFP signal. The scale bars are 5 μm.

**Figure S6.** Parental Cf-C1 lack GFP signal by confocal and western blot analyses. Deconvolved confocal images of GFP and DAPI images of fixed swimmer and attached cell cultures taken with the same settings as shown in Figure 6. Fluorescent images are shown individually and merged. A brightfield image for each is included for reference and was used to generate the dashed outline of the flagellum in the merged image. A western blot of extracts of swimming and attached cells probed with anti-GFP and anti-BiP (loading control) antibodies.

**Video S1.** 4×10^5^ parental Cf-C1 cells were added to wells of a 24-well plate and imaged at 75 second intervals for 16 hours to obtain movies of *C. fasciculata* as they adhere. This video shows the full imaging frame at 60x magnification, which highlights several adhesion events occurring during the imaging time. Note an oblong out-of-focus object appears to the left of center. The scale bar is 50 μm.

**Video S2.** Close up and cropped version of Video S1. For reference, the same oblong out-of-focus object that is visible near the center of Video S1 appears at the left edge of this video. Note when a dividing pair of cells first adheres to the culture dish surface (time 5:42:30), one cell swims away. The other remains characteristically attached at the junction between the cell body and the flagellum. Both the cell body and the flagellum move independently until the flagellum is retracted and the cell body is round (time 8:40:00). The first division as an adherent cell occurs at approximately 9:08:45. One of the adherent daughters divides again approximately 4 hours and 4 minutes later (time 13:12:30). The other adherent daughter divides approximately 5 hours and 25 minutes later (time 14:33:45). The scale bar is 10 μm.

## Notes

### Competing Interest Statement

The authors have declared no competing interest.

### Summary of Updates

Addition of supplemental materials

