## Supplemental table 1 for "Adhesion of *Crithidia fasciculata* promotes a rapid change in developmental fate driven by cAMP signaling"

| Primer name | Sequence |
|---|---|
| pLENTeYFP_f | 5'-gtgagcaagggcgaggagc-3' |
| pLENTNeo_r | 5'-tcagaagaactcgtcaagaag-3' |
| PF16cterm500_f | 5'-aaggaggcggaggatcata-3' |
| PF16cterm_r | 5'-gctcctcgcccttgctcacgtgctgctgcacgtggtagttc-3' |
| PF16utr_f | 5'-cttcttgacgagttcttctgaaaaaaaaagggaaggagaggcgac-3' |
| PF16utr500_r | 5'-atagtgatgcttgccgtcgt-3' |
| PF16_500bpnest_f | 5'-gctgctgcctggtcact-3' |
| PF16_500bpnest_r | 5'-ccttccttcactgcctctgt-3' |
| PF16ORF_f | 5’-gacgatccagcgtatcaagg-3’ |
| PF16UTR_r | 5’-tctttgcattgagcgagcta-3’ |
| CfRAC6500_Nd_f | 5’-gcggcgcatatgatgtccccccctctgtggacgc-3’ |
| CfRAC6500_Kp_r | 5’-gcggcgggtaccctgcttgcccgaaaatccatt-3’ |
| CfPDEA_Nd_f | 5’-tatcatatgatgtccgatttcaaagagcag-3' |
| CfPDEA_Bg_r | 5’-tatagatctcgagtcatcgtggctagcccag-3' |
