## Supplementary figures and images for "Adhesion of *Crithidia fasciculata* promotes a rapid change in developmental fate driven by cAMP signaling"

### Figure S1

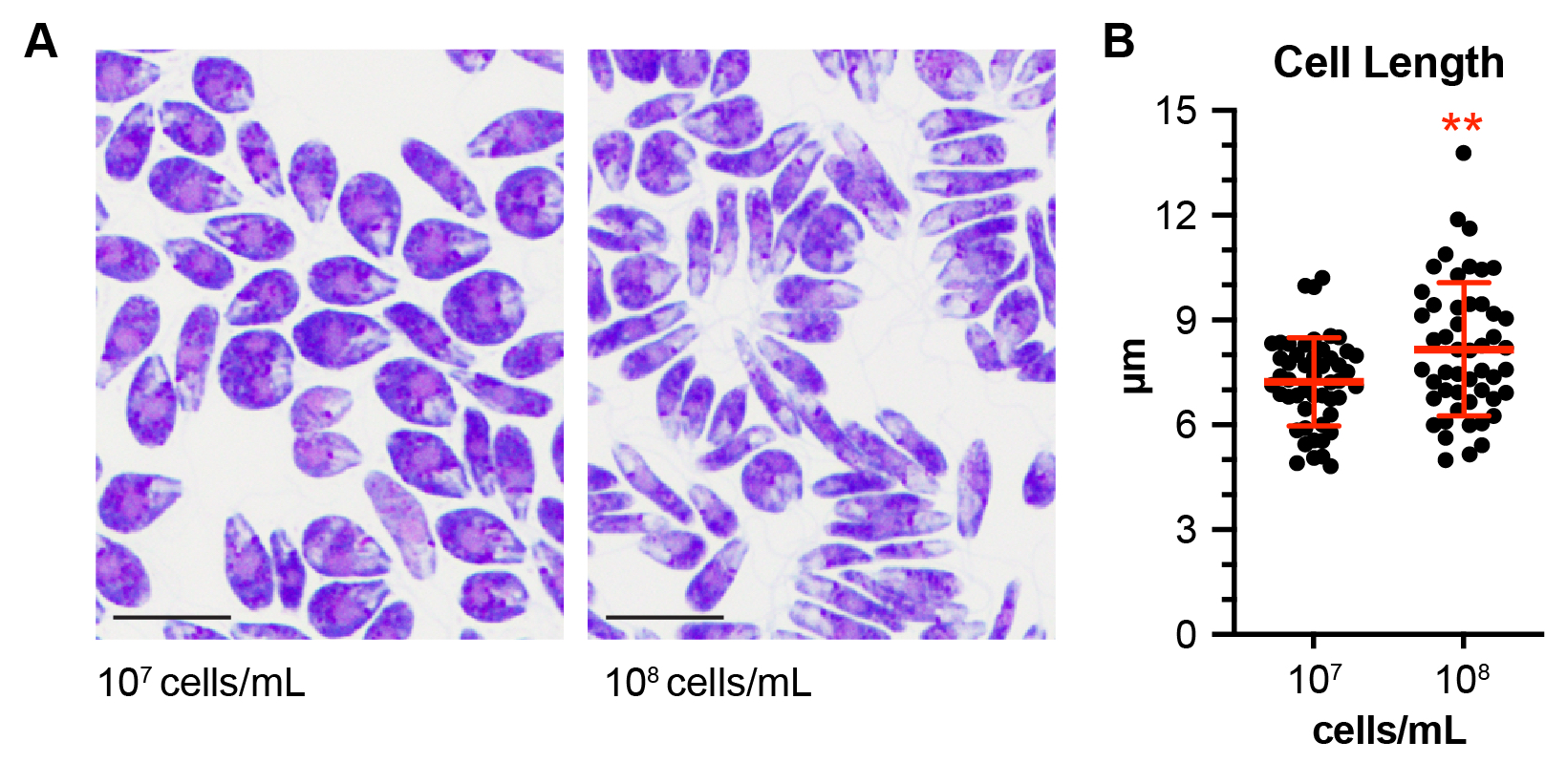

### Figure S2

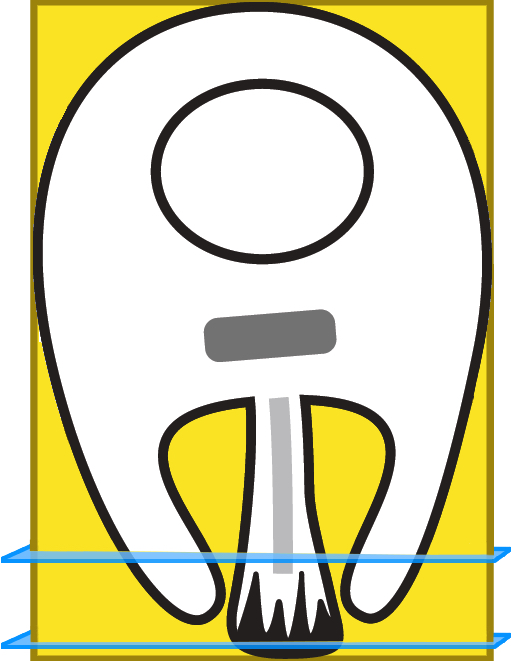

### Figure S3

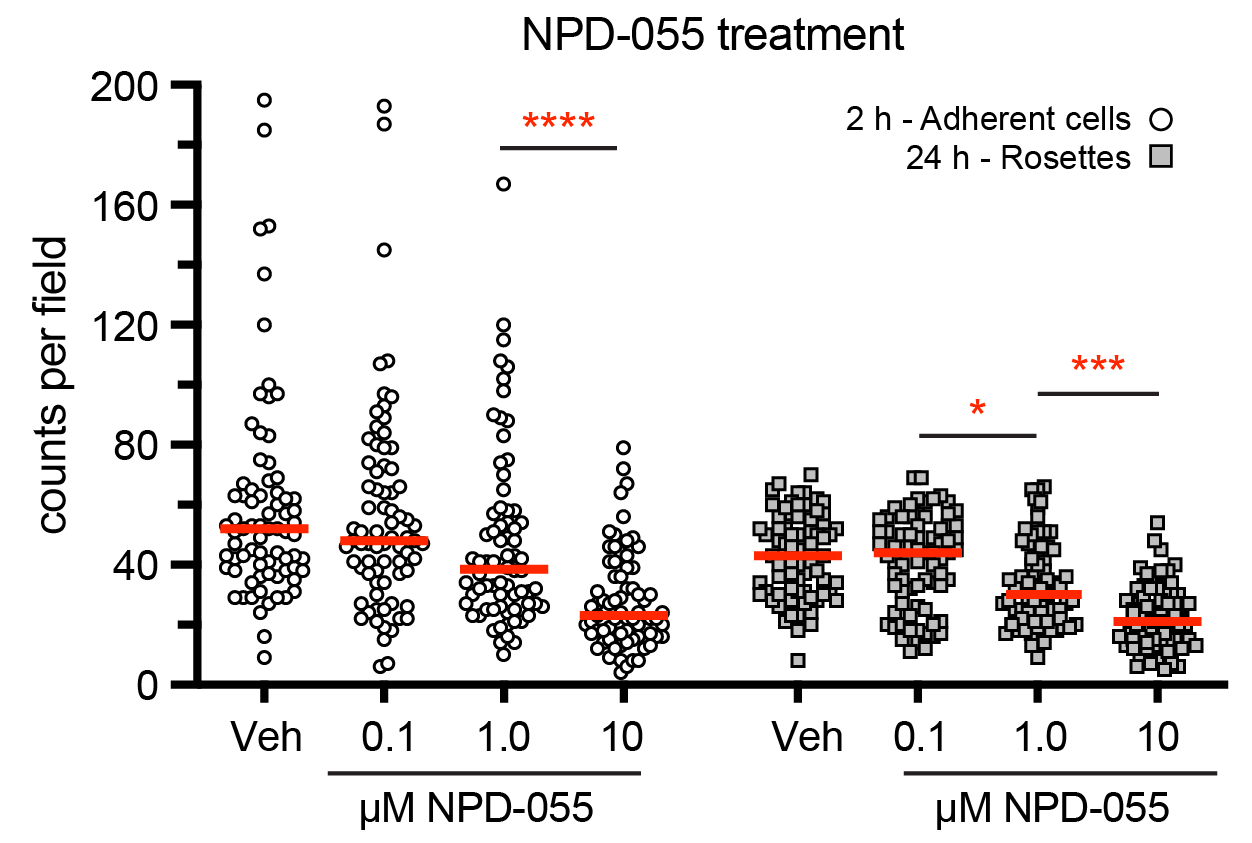

### Figure S4

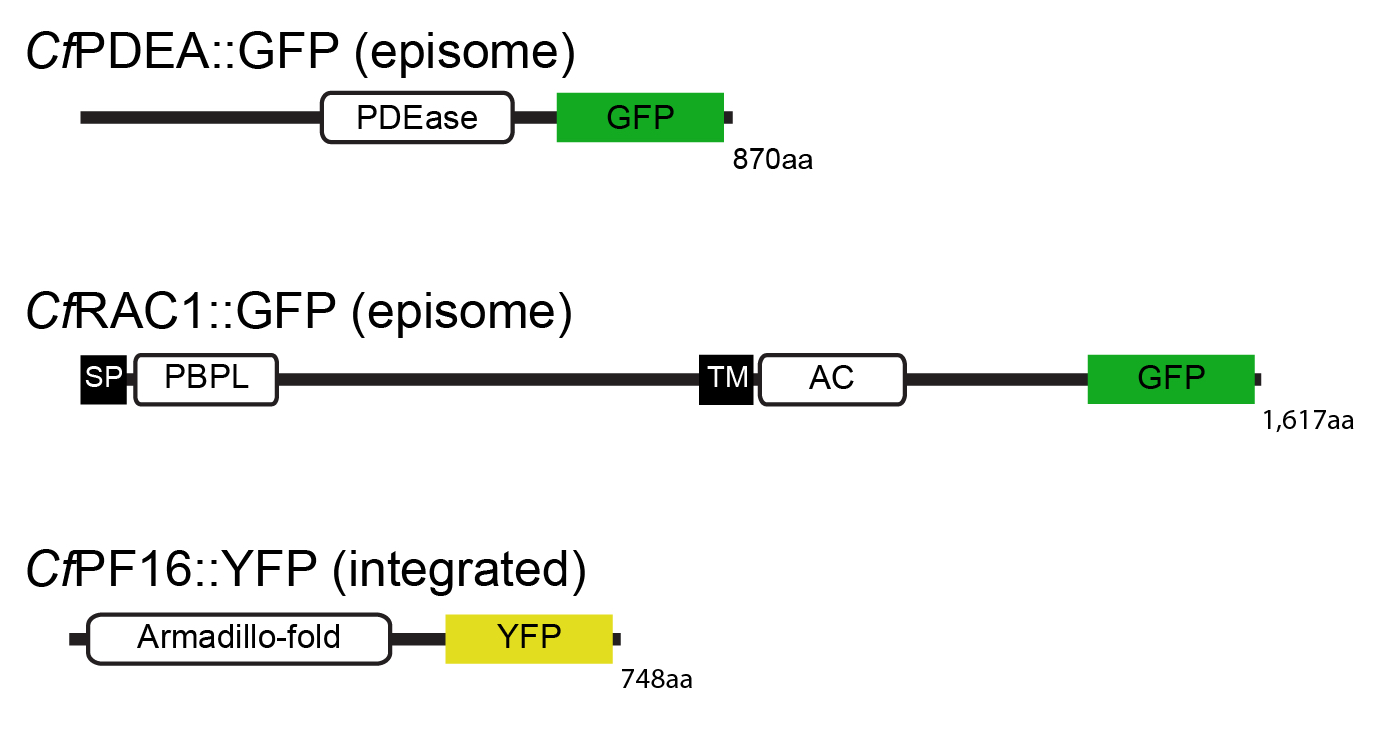

### Figure S5

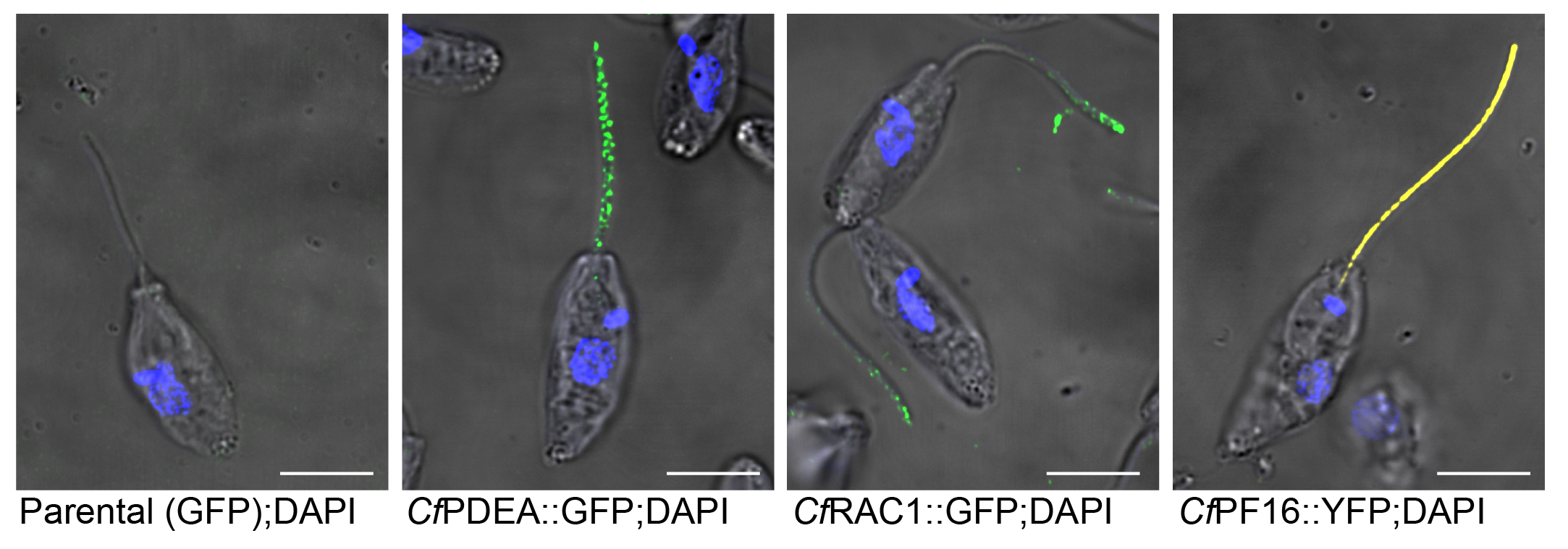

### Figure S6

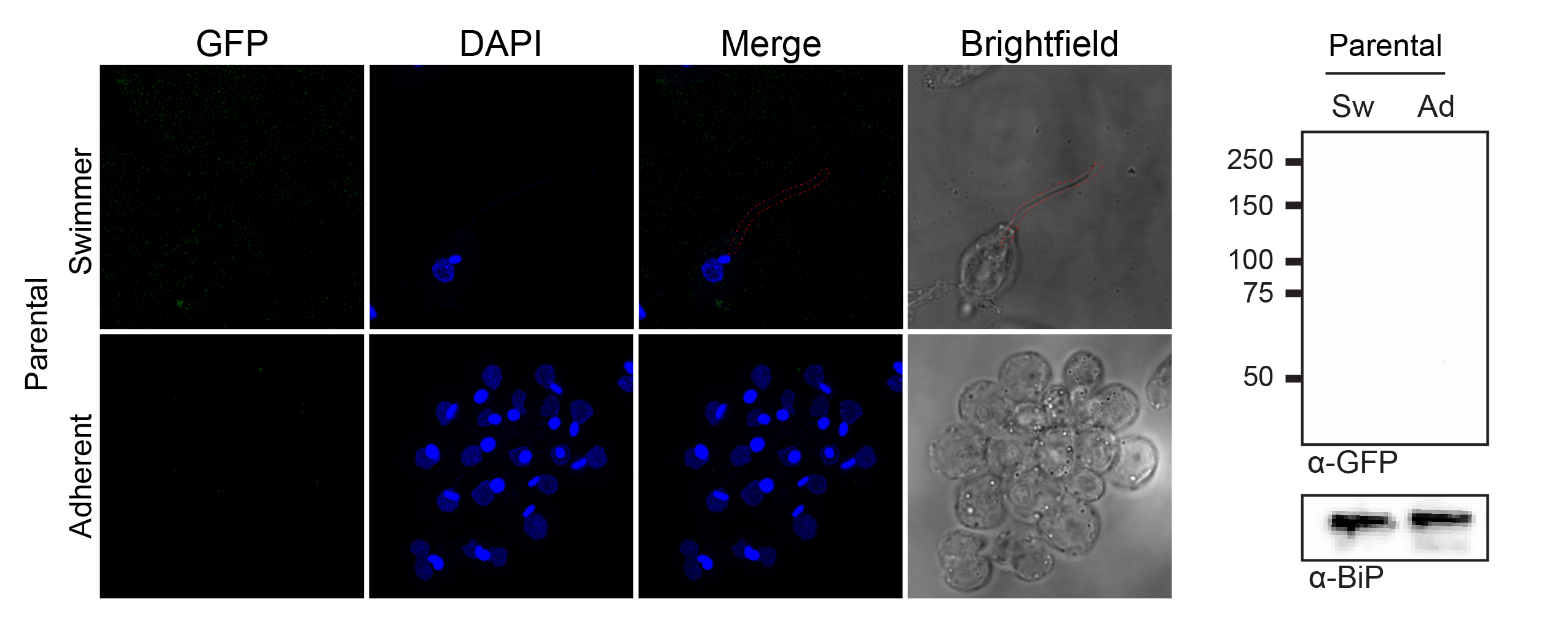

### Striking Image

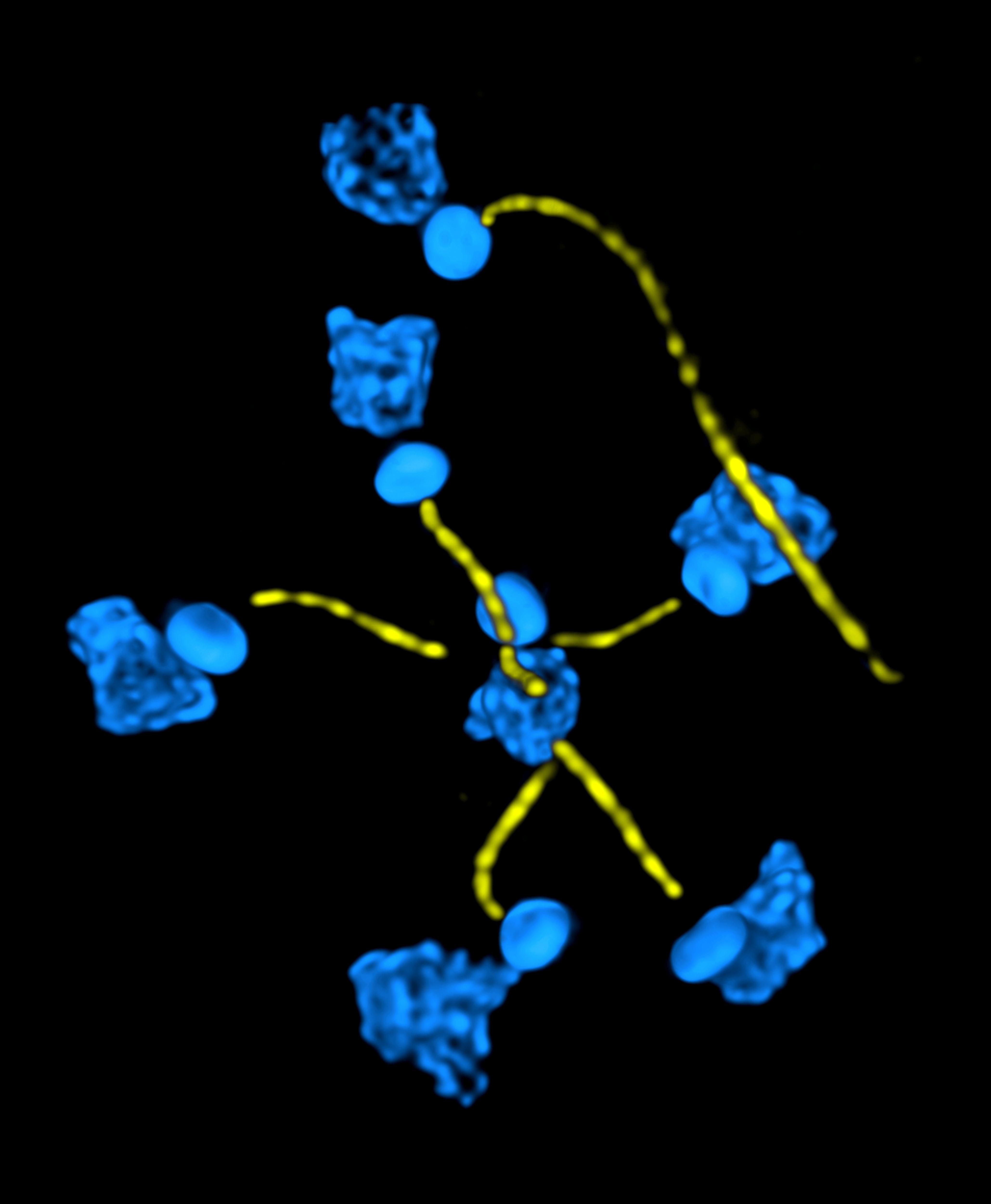
